# RADICAL CATION SCAVENGING ACTIVITY OF BERBERINE BRIDGE ENZYME-LIKE OLIGOSACCHARIDE OXIDASES ACTING ON SHORT CELL WALL FRAGMENTS

**DOI:** 10.1101/2022.11.03.515065

**Authors:** Anna Scortica, Valentina Scafati, Moira Giovannoni, Manuel Benedetti, Benedetta Mattei

**Affiliations:** Department of Life, Health and Environmental Sciences, University of L’Aquila, 67100 L’Aquila, Italy

**Author notes:** correspondence, Department of Life, Health and Environmental Sciences, University of L’Aquila, 67100 L’Aquila, Italy. These authors equally contributed to this work (Anna Scortica, Valentina Scafati, Moira Giovannoni).

## Abstract

Oligogalacturonide-oxidases (OGOXs) and cellodextrin-oxidase (CELLOX) are plant berberine bridge enzyme-like oligosaccharide-oxidases (OSOXs) that oxidize, respectively, oligogalacturonides (OGs) and cellodextrins (CDs), thereby inactivating their elicitor nature and concomitantly releasing H_2_O_2_. Little is known about the physiological role of OSOX activity. By using an ABTS^•+^-reduction assay, we identified a novel reaction mechanism through which the activity of OSOXs on cell wall oligosaccharides scavenged the radical cation ABTS^•+^ with an efficiency dependent on the type and length of the oxidized oligosaccharide. In contrast to the oxidation of longer oligomers such as OGs (degree of polymerization from 10 to 15), the activity of OSOXs on short galacturonan- and cellulose-oligomers (degree of polymerization ≤ 4) successfully counteracted the radical cation-generating activity of a fungal laccase, suggesting that OSOXs can generate radical cation scavenging activity in the apoplast with a power proportional to the extent of degradation of plant cell wall, with possible implications for redox homeostasis and defense against oxidative stress.

## INTRODUCTION

Plants are constantly menaced by a wide array of pathogens such as virus, bacteria and fungi. Against them, plants evolved a robust barrier composed of polysaccharides, phenols and proteins, i.e., the cell wall, and a sophisticated defence system that can be promptly activated at the occurrence. Phytopathogenic fungi decompose the cell wall by the action of several cell wall degrading enzymes (CWDEs), a wide category of enzymes that includes glycoside hydrolases (GHs), lyases and oxidoreductases^1,2^. The enzymatic hydrolysis of cell wall polysaccharides results in the initial accumulation of cell wall fragments that can be perceived by plants as danger signals^3^; as the pathogen attack continues, these cell wall fragments are progressively converted into dimers and monomers that microbes use as carbon source to sustain their infection process^1,4^. The identification of different berberine bridge enzyme-like (BBE-l) proteins from *Arabidopsis thaliana* as specific oligosaccharide oxidases (OSOXs) paved the way to novel insights in plant immunity. To date, BBE-l proteins acting as true OSOXs are oligogalacturonide-oxidases 1-4 (OGOX1-4) and cellodextrin-oxidase (CELLOX). The four OGOX isoforms and CELLOX are capable of oxidizing, respectively, oligogalacturonides (OGs) and cellodextrins (CDs), thereby inactivating their elicitor nature and concomitantly releasing H_2_O_2_, a molecule with multiple physiological roles^5^. Besides an increased recalcitrance to enzymatic hydrolysis displayed by the oxidized oligosaccharides^6^, nothing is known about their involvement in other physiological processes. Recently, 2,2’-azino-bis (3-ethylbenzothiazoline-6-sulfonic acid) (ABTS)-peroxidase (POD) based assays were successfully used to measure the activity of fungal oligosaccharide oxidases^7^ and Arabidopsis OGOX1^8^. In the latter study, the combined use of OGOX1 and a horseradish POD (HRP VI-A type) allowed the measurement of the OG-oxidizing activity in continuous mode. This allowed, in turn, to monitor *in vitro* the capability of OSOXs of converting the activity of microbial GHs into a controlled H_2_O_2_-dependent oxidative signal, by setting up an enzymatic machinery composed of a microbial GH, a plant OSOX and a plant POD^9^. The main requirement for the functioning of such machinery was that the GH/OSOX pair shared the same substrate specificity, i.e., capability of hydrolysing/oxidizing the same cell wall carbohydrate. By using an ABTS^•+^-reduction assay, we identified here a novel reaction mechanism through which the activity of OSOXs on cell wall oligosaccharides scavenged the radical cation ABTS^•+^ with an efficiency dependent on the type and length of the oxidized oligosaccharides. In contrast to the oxidation of longer cell wall oligosaccharides (degree of polymerization from 10 to 15), the activity of OSOXs on short cell wall oligomers (degree of polymerization ≤ 4) successfully counteracted the radical cation-generating activity of a fungal laccase, suggesting that OSOXs can generate radical cation scavenging activity in the apoplast with a power proportional to the extent of degradation of plant cell wall.

## RESULTS

### Heterologous expression of FHS-OSOXs in *P. pastoris*

The flag-his-sumoylated form of CELLOX (FHS-CELLOX) was obtained as previously described^9^. The sequence encoding the flag-his-sumoylated form of OGOX1, here referred to as FHS-OGOX1, was cloned under the control of the methanol-inducible promoter AOX, expressed in *P. pastoris* and purified in a single step by IMAC chromatography. The purity grade of FHS-OGOX1 preparation was assessed by SDS-PAGE/Coomassie blue staining analysis (Fig. S1a) whereas the protein concentration was determined by UV-absorbance. In accordance with this procedure, the final protein yield from the highest expressing Pichia transformant ranged from 2 to 4 mg.L^−1^. The specific activity of FHS-OGOX1 on OGs and two different galacturonan oligomers, i.e., tri-galacturonic acid (OG3) and tetra-galacturonic acid (OG4), is summarized in Fig. S1b. It is worth noting that all the enzymatic assays described here were performed at apoplastic pH value (5.0) and by using an oligosaccharide concentration (15 μM) compatible with that used to trigger the defense responses in plants^8^.

### The activity of OSOXs on short oligosaccharides scavenged the radical cation ABTS^•+^

To date, the activity of OSOXs can be evaluated by two different methods, i.e., the xylenol orange assay^6,10^ and the ABTS-HRP coupled assay^8^ (Fig. 1a). Differently from the xylenol orange assay, the ABTS-HRP coupled assay was unable to detect the activity of FHS-OGOX1 on OG4 (Fig. 1b). To deeper investigate this surprising result, we modified the buffer composition of the ABTS-HRP coupled assay by using chemically oxidized ABTS^•+^ as substrate. In brief, the starting amount of ABTS was converted into 82% ABTS^•+^ and 18% ABTS by using K_2_S_2_O_8_ as ABTS-activator and, importantly, HRP was excluded from the assay, thus reducing the number of experimental variables of the enzymatic reaction. Moreover, the concomitant presence of ABTS^•+^ and ABTS in the buffer allowed to identify all the ongoing redox reactions of the ABTS/ABTS^•+^ pair that, otherwise, could not be clearly detected in the presence of only one species (Fig. 1b). The use of such assay, hereafter referred to as ABTS^•+^-reduction assay, revealed that the activity of FHS-OGOX1 on OG4, and at lesser extent on OG3 and OGs, decreased the amount of radical cation ABTS^•+^ over reaction time (Fig. 2a). This result indicated that the lack of oxidizing activity of FHS-OGOX1 on OG4 as revealed by the ABTS-HRP coupled assay (Fig. 1b) was due to a side-reaction consisting in a time-dependent scavenging of ABTS^•+^ (Fig. 2a). In parallel, the amount of H_2_O_2_ as produced by FHS-OGOX1 was evaluated under the same reaction conditions (i.e., same pH value and enzyme/substrate concentration) by using the xylenol orange assay (Fig. 2b). A time-dependent reduction of ABTS^•+^ occurred also in the presence of FHS-CELLOX and two different CD-oligomers, i.e., cellotetraose (CD4) and cellotriose (CD3) (Fig. 3a). Also in this case, the amount of H_2_O_2_ as produced by FHS-CELLOX was evaluated under the same reaction conditions (i.e., same pH value and enzyme/substrate concentration) by using the xylenol orange assay (Fig. 3b). Although OGOX1 and CELLOX are characterized by different enzymatic properties^6,9,10^, the ABTS^•+^-reduction assay was set on concentrations of enzymes and substrates that produced similar rates of H_2_O_2_ (Fig. 2b-3b), making easier to compare the ABTS^•+^-scavenging activity of the two FHS-OSOXs.

**Figure 1.**
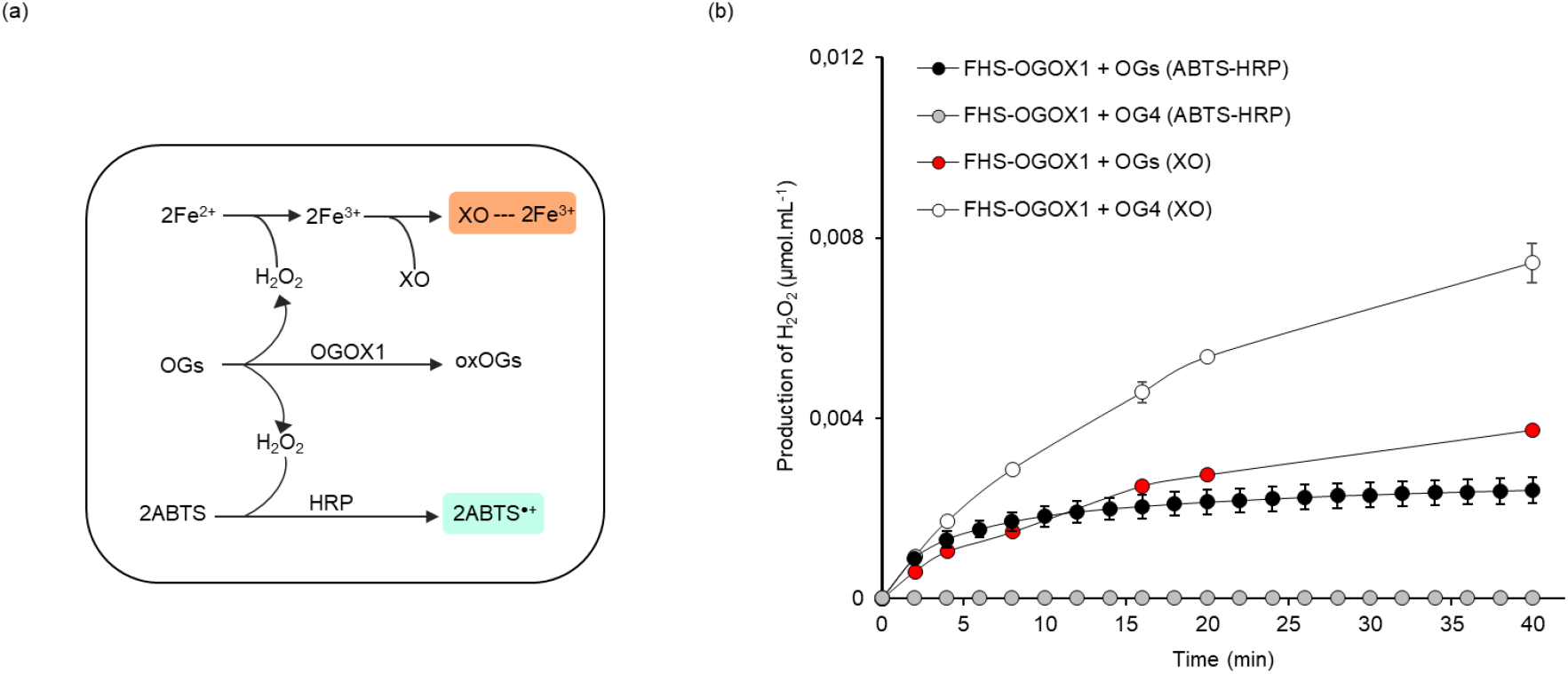
HRP-mediated oxidation of ABTS is not observed in the presence of FHS-OGOX1 and OG4. (a) Schematic representation showing the reactions of (up) the xylenol orange (XO) and (down) ABTS-HRP coupled assays as used for the kinetic measurements of FHS-OGOX1 using OGs as substrate. (b) Oxidizing activity of FHS-OGOX1(μmoles of H_2_O_2_.mL^−1^) at pH 5.0 using equimolar amounts of OGs and OG4 by xylenol orange (XO) and ABTS-HRP coupled (ABTS-HRP) assay. Values are mean ± SD (N=3). [HRP: horseradish peroxidase, OGs: oligogalacturonides, OG4: tetra-galacturonic acid, OGOX1: oligogalacturonide-oxidase 1 from *A. thaliana*, XO: xylenol orange, oxOGs: oxidized OGs].

**Figure 2.**
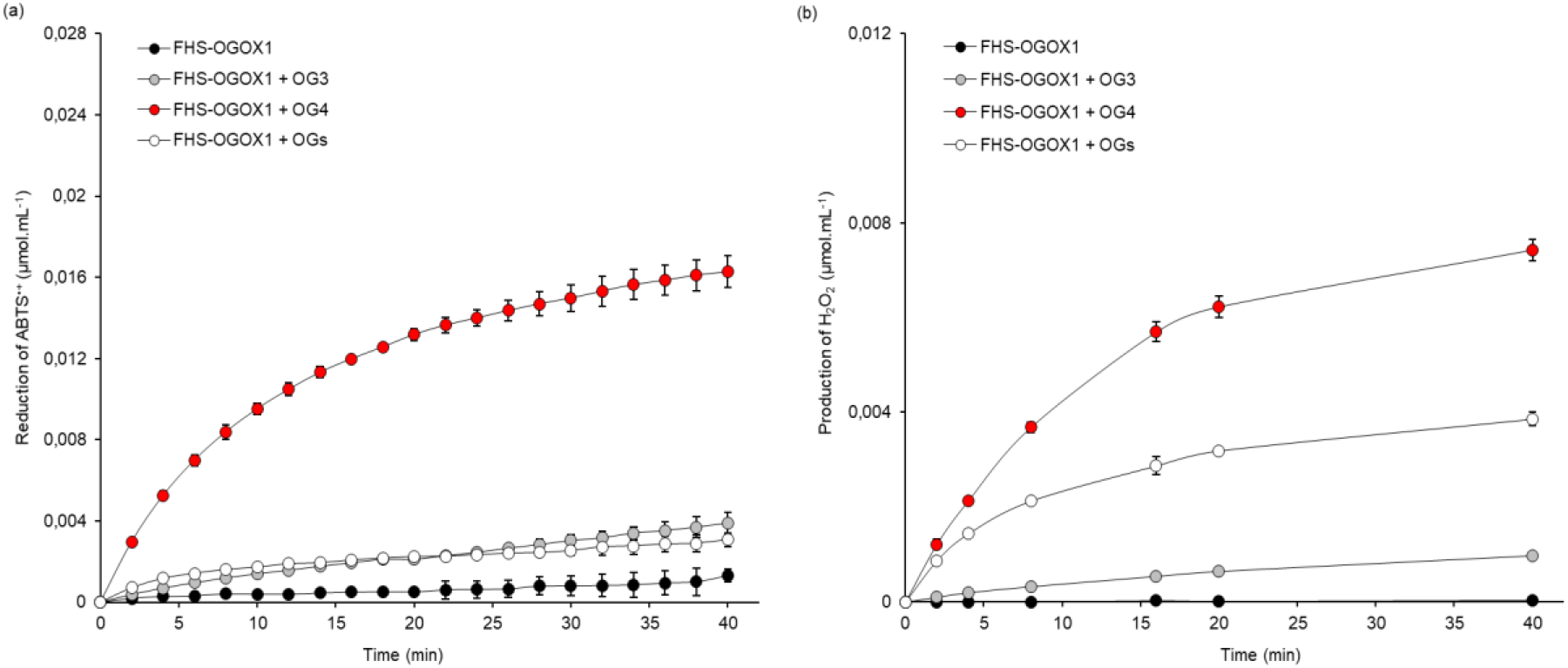
Scavenging activity of FHS-OGOX1 and short OG-oligomers towards the radical cation ABTS^•+^. (a) Reduction of ABTS^•+^ (μmol.mL^−1^) over time at pH 5.0 by the activity of FHS-OGOX1 using equimolar amounts of OG3, OG4 and OGs. (b) Production of H_2_O_2_ (μmol.mL^−1^) over time by FHS-OGOX1 under the same reaction conditions (same pH value and enzyme/substrate concentration) shown in (a). Values are mean ± SD (N=3). [FHS-OGOX1: flag-his-sumoylated oligogalacturonide-oxidase 1, OGs: oligogalacturonides, OG4: tetra-galacturonic acid, OG3: tri-galacturonic acid].

**Figure 3.**
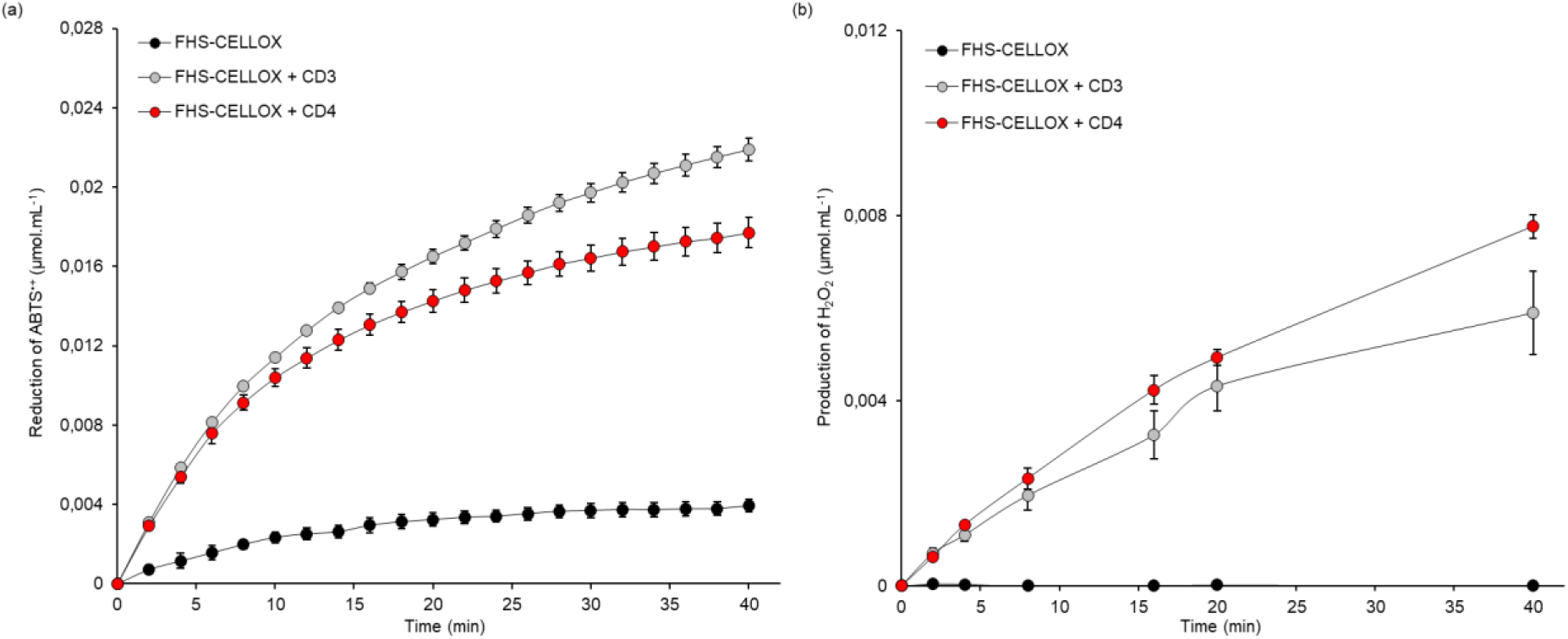
Scavenging activity of FHS-CELLOX and short CD-oligomers towards the radical cation ABTS•^+^. (a) Reduction of ABTS^•+^ (μmol.mL^−1^) over time at pH 5.0 by the activity of FHS-CELLOX using equimolar amounts of CD3 and CD4. (b) Production of H_2_O_2_ (μmol.mL^−1^) over time by FHS-CELLOX under the same reaction conditions (same pH value and enzyme/substrate concentration) shown in (a). Values are mean ± SD (N=3). [CD4: cellotetraose, CD3: cellotriose, FHS-CELLOX: flag-his-sumoylated cellodextrin-oxidase].

### The scavenging efficiency of different OSOX/oligomer combinations is dependent on the type and length of each oligomer

To assess the scavenging efficiency of different FHS-OSOX/oligomer combinations, the net reduction of ABTS^•+^ as obtained by the activity of OSOXs on the vary oligosaccharides (Fig. 2a-3a) was normalized to the amount of H_2_O_2_ as produced under the same reaction conditions (Fig. 2b-3b). These results are summarized in Table I. The scavenging efficiency of different FHS-OSOX/oligomer combinations towards ABTS^•+^ was inversely proportional to the length of each oligosaccharide (Table I). In particular, the activity of FHS-OGOX1 scavenged 2.5 moles of ABTS^•+^ per mole of oxidized OG3, about 2.1 moles of ABTS^•+^ per mole of oxidized OG4, and 0.6 mole of ABTS^•+^ per mole of oxidized OGs (Table I). A higher radical cation scavenging activity was observed for both FHS-CELLOX/CD-oligomer combinations since the activity of FHS-CELLOX scavenged 4 and 2.9 moles of ABTS^•+^ per mole of oxidized CD3 and CD4, respectively (Table I). Also in this case, the activity of FHS-CELLOX on the shortest cellodextrin (CD3) produced the highest scavenging activity towards ABTS^•+^ (Table I). In accordance with the more oxidized state of galacturonan with respect to cellulose, the scavenging efficiency of FHS-OGOX1 was lower than that of FHS-CELLOX for oligosaccharides with the same length (Table I).

**Table I.** Scavenging efficiency of different OSOX/oligomer combinations towards the radical cation ABTS^•+^. Scavenging efficiency of different OSOX/oligomer combinations towards the radical cation ABTS^•+^ as determined by the ABTS^•+^-reduction and xylenol-orange assays. Values “a,b” were extrapolated from the kinetic analysis shown in Fig. 2-3 at three different reaction times (4, 8 and 16 or 20 min). For each FHS-OSOX/oligomer combination, the value of scavenging efficiency was calculated as mean ± SD (N=3). [CD4: cellotetraose, CD3: cellotriose, FHS-CELLOX: flag-his-sumoylated cellodextrin-oxidase, FHS-OGOX1: flag-his-sumoylated oligogalacturonide-oxidase 1, OGs: oligogalacturonides, OG4: tetra-galacturonic acid, OG3: tri-galacturonic acid, ox-oligomer: oxidized oligomer].

| Reaction time (min) | FHS-OSOX/oligomer | a, Net reduction of ABTS<sup>•+</sup> ($\mu\text{mol} \cdot \text{mL}^{-1}$) | b, Production of H<sub>2</sub>O<sub>2</sub>/ox-oligomer ($\mu\text{mol} \cdot \text{mL}^{-1}$) | a.b<sup>-1</sup>, Scavenging efficiency (ABTS<sup>•+</sup>.ox-oligomer<sup>-1</sup>) |
| --- | --- | --- | --- | --- |
| 4 | FHS-OGOX1/OGs | 0,0009 ± 7,6E<sup>-05</sup> | 0,0014 ± 3,4E<sup>-05</sup> | 0,6 ± 0,04 |
| 8 | | 0,0012 ± 0,0001 | 0,0021 ± 5,7E<sup>-05</sup> | |
| 16 | | 0,0016 ± 0,0002 | 0,0029 ± 0,0002 | |
| 4 | FHS-OGOX1/OG4 | 0,0049 ± 0,0002 | 0,0021 ± 4,5E<sup>-05</sup> | 2,1 ± 0,1 |
| 8 | | 0,0079 ± 0,0003 | 0,0037 ± 0,0001 | |
| 16 | | 0,0114 ± 0,0001 | 0,0057 ± 0,0002 | |
| 8 | FHS-OGOX1/OG3 | 0,0008 ± 7,8E<sup>-05</sup> | 0,0003 ± 2,4E<sup>-05</sup> | 2,5 ± 0,1 |
| 16 | | 0,0014 ± 0,0001 | 0,0005 ± 2,3E<sup>-05</sup> | |
| 20 | | 0,0016 ± 0,0001 | 0,0006 ± 4,5E<sup>-05</sup> | |
| 4 | FHS-CELLOX/CD4 | 0,0042 ± 0,0003 | 0,0013 ± 6,3E<sup>-05</sup> | 2,9 ± 0,4 |
| 8 | | 0,0071 ± 0,0004 | 0,0023 ± 0,0002 | |
| 16 | | 0,0101 ± 0,0005 | 0,0042 ± 0,0003 | |
| 4 | FHS-CELLOX/CD3 | 0,0047 ± 1,3E<sup>-05</sup> | 0,0011 ± 0,0001 | 4 ± 0,3 |
| 8 | | 0,0079 ± 0,0001 | 0,0019 ± 0,0003 | |
| 16 | | 0,0119 ± 0,0002 | 0,0033 ± 0,0005 | |

### The radical cation scavenging activity towards ABTS^•+^ requires an ongoing activity of OSOX on short oligosaccharides

ABTS is also used as substrate to evaluate the Trolox Equivalent Antioxidant Capacity (TEAC) of several antioxidant molecules^11^. Since the reaction catalyzed by OSOXs is highly complex^3,6,8^ and different by-products could be involved, additional experiments were performed in order to better elucidate the reaction mechanism by using FHS-OGOX1/OG4 as representative FHS-OSOX/oligomer combination (Fig. S2). The absence of a clear scavenging effect in the presence of FHS-OGOX1, H_2_O_2_, OG4, [H_2_O_2_ + OG4] or [H_2_O_2_ + oxidized OG4 (oxOG4)] demonstrated that an ongoing activity of FHS-OGOX1 on OG4 is essential for the time-dependent reduction of ABTS^•+^ (Fig. S2). Unfortunately, the ABTS^•+^-reduction assay contains an amount of ABTS^•+^ (0.09 μmol.mL^−1^) that interferes with the xylenol orange, impeding its utilization for a direct measurement of H_2_O_2_ during the reaction of scavenging. To detect any eventual production of H_2_O_2_, three different FHS-OGOX1/oligomer combinations were assayed in the presence of an excess of HRP, here used to counter-oxidize the scavenged ABTS^•+^ by consuming the H_2_O_2_ produced from the different OSOX/oligomer combinations (Fig. 1a)^8^. In the presence of HRP, the ABTS^•+^-scavenging activity of both FHS-OGOX1/OG3 and FHS-OGOX1/OG4 combinations remained almost the same, whereas an opposite activity (production of ABTS^•+^) was observed for the FHS-OGOX1/OGs pair (Fig. S3). Based on this result (Fig. S3), the presence of H_2_O_2_ was only detected in the reaction containing FHS-OGOX1/OGs, i.e., the combination with the lowest scavenging efficiency (Table I).

### The ABTS-oxidizing activity of a fungal laccase is counteracted by the activity of OSOXs on short oligosaccharides

To evaluate whether the scavenging activity of OSOXs can counteract the oxidizing activity of microbial ligninases, the laccase from *Trametes versicolor* was assayed in the presence of different FHS-OSOX/oligomer combinations by using ABTS as substrate. It is worth noting that the oxidation of ABTS by laccase, differently from HRP, is not H_2_O_2_-dependent. In accordance with the scavenging efficiencies reported in Table I, the ABTS-oxidizing activity of laccase was reduced with the highest efficiency by the FHS-CELLOX/CD3 combination and with lower efficiency by the FHS-CELLOX/CD4 and FHS-OGOX1/OG4 combinations, whereas the FHS-OGOX1/OGs combination was ineffective (Fig. 4). Indeed, this result clearly demonstrated that the scavenging activity of OSOXs towards the laccase-catalyzed oxidation of ABTS required the presence of oxidizable short oligosaccharides. Differently from the ABTS^•+^-reduction assay, the lower amount of ABTS^•+^ (0-0.009 μmol.mL^−1^, Fig. 4) as contained in the ABTS-oxidation assay allowed a direct measurement of H_2_O_2_ by using the xylenol orange. In the presence of laccase, the (negligible) scavenging activity of the FHS-OGOX1/OGs combination did not have any impact on the production of H_2_O_2_ whereas the strong scavenging activities of both FHS-OGOX1/OG4 and FHS-CELLOX/CD3 combinations were associated with decreased levels of H_2_O_2_ (Fig. S4). Notably, the reduced amounts of H_2_O_2_ from the activity of FHS-OGOX1/OG4 and FHS-CELLOX/CD3 combinations (0.003 μmol.mL^−1^ and 0.004 μmol.mL^−1^) were compatible with the amounts of scavenged ABTS^•+^ (0.005 μmol.mL^−1^ and 0.008 μmol.mL^−1^) (Fig. S4). Also in this case, higher scavenging efficiencies were related to lower levels of H_2_O_2_ and vice versa (Table I).

**Figure 4.**
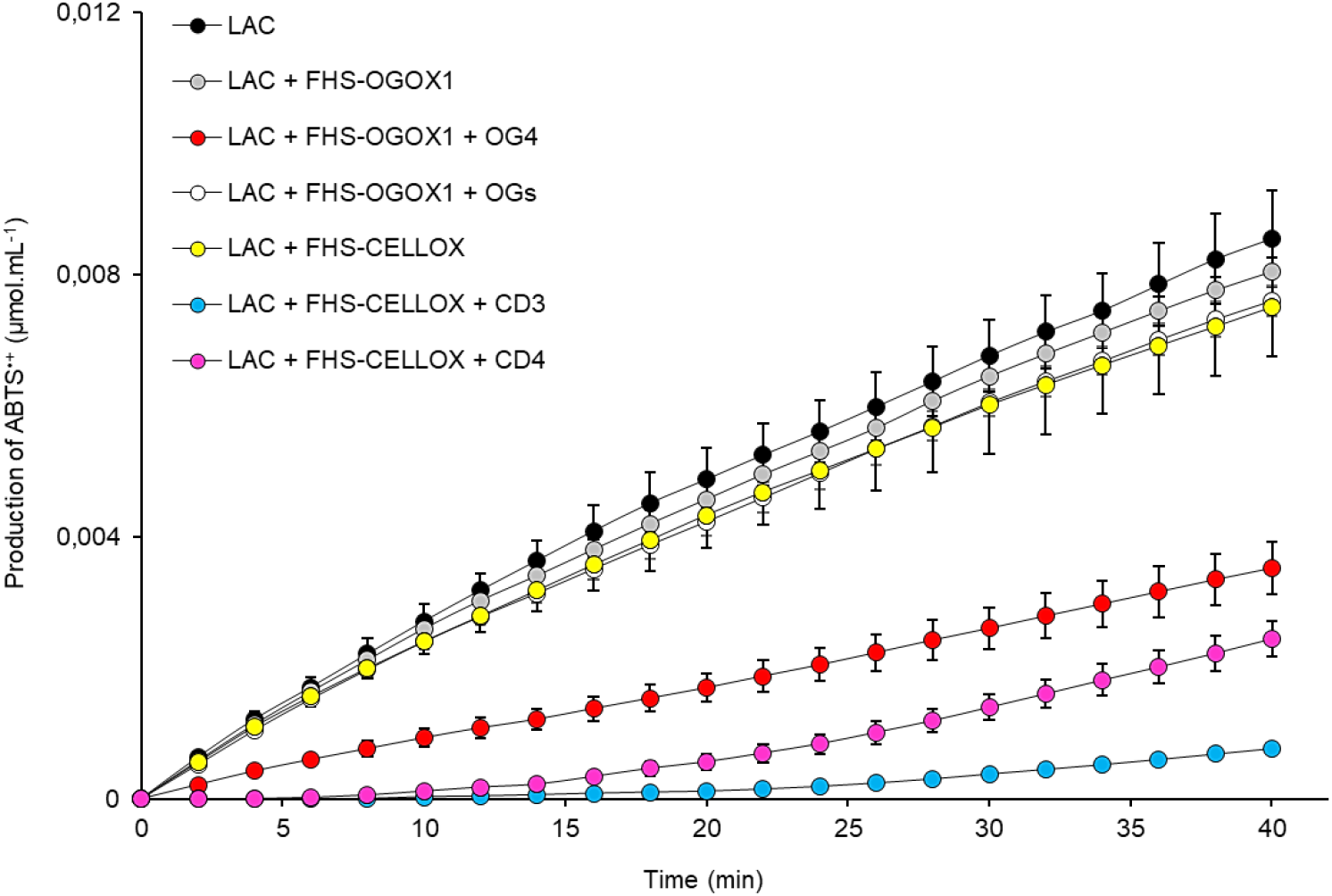
The activity of FHS-OSOXs on short oligosaccharides counteracts the laccase-catalyzed oxidation of ABTS. Production of ABTS^•+^ (μmol.mL^−1^) over time at pH 5.0 by laccase from *T. versicolor* alone and in the presence of FHS-OGOX1 and two different galacturonic acid oligomers (OG4 and OGs), and of FHS-CELLOX and two different cellodextrins (CD3 and CD4). Laccase activity was not affected in the only presence of oligomers (not shown). Values are mean ± SD (N=3). [CD3: cellotriose, CD4: cellotetraose, FHS-CELLOX: flag-his-sumoylated cellodextrin-oxidase, FHS-OGOX1: flag-his-sumoylated oligogalacturonide-oxidase 1, LAC: laccase from *T. versicolor*, OGs: oligogalacturonides, OG4: tetra-galacturonic acid].

In conclusion, the OSOX activity on short oligosaccharides scavenged the radical cation ABTS^•+^ (Fig. 2-4, Fig. S2) with an efficiency dependent on the composition and length of each oligosaccharide (Table I) and at the expense of the H_2_O_2_ as produced from their oxidation (Fig. S3-S4).

## DISCUSSION

POD-mediated oxidation of ABTS is well-documented since 1975^12^; subsequently, the use of ABTS was also extended to fungal laccases as redox mediator in lignin depolymerization^13^. To our knowledge, this is the first report describing a clear enzyme-dependent scavenging activity towards the radical cation ABTS^•+^. Although the use of ABTS as mediator in biological redox reactions^14^ can be criticized due to its synthetic nature, the scavenging activity towards ABTS^•+^ was achieved here by using nanomolar concentration of OSOXs (4-16 nM) and micromolar concentration of each oligosaccharide (15 μM); notably, the latter substrate concentration was about 150-3000 folds lower than that reported for the non-enzymatically catalyzed ABTS^•+^-reduction reactions^15^. Moreover, the radical cation scavenging activity was generated by OSOXs with different substrate specificities and was markedly dependent on the type and length of oxidized oligosaccharides, indicating possible distinct involvement in the plant immune response (Fig. 1-4). *In vivo*, the scavenging activity of different OSOX/oligomer combinations may be directed towards metal cofactors, lignol radicals, oxidized antioxidants and/or unknown molecules of plant and of microbial origin with redox potentials similar to the radical cation ABTS^•+^ (Fig. 5a).

**Figure 5.**
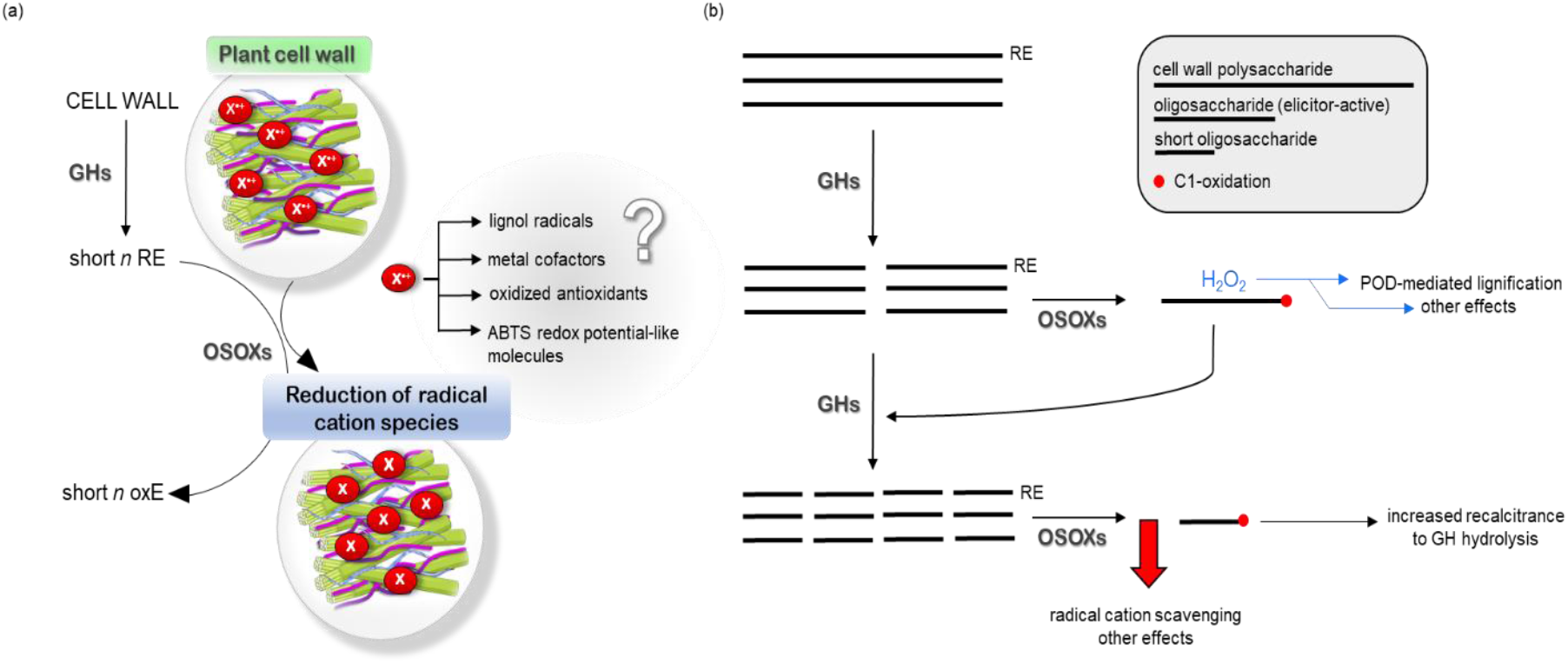
Proposed models of OSOXs as cell wall defense proteins. (a) Scavenging of unknown radical cations (X^•+^) by the activity of OSOXs on short cell wall oligosaccharides. (b) Potential contribution of reaction products from OSOX activity in the maintenance of cell wall integrity during the infection. [GH: glycoside hydrolase, OSOX: berberine bridge enzyme-like oligosaccharide oxidase, oxE: oxidized end, POD: plant peroxidase, RE: reducing end].

To date, OGOX1-4 and CELLOX are the only plant BBE-l enzymes with proven oxidizing activities towards cell wall fragments with elicitor nature, i.e., OGs and CDs, and therefore they can be classified as true OSOXs. The physiological role of OSOXs is still under investigation. During the degradation of plant cell wall, the resulting cell wall fragments can be converted by OSOXs into H_2_O_2_ and oxidized oligosaccharides. By using a multiple enzyme-based assay, we have already demonstrated that H_2_O_2_ from OSOX activities may be used by extracellular PODs to reinforce the cell wall in a manner proportional to the entity of cell wall damage taken^9^. Here we found a possible involvement of OSOX activity on short cell wall oligomers as scavenger towards the radical cation-producing activity of fungal laccases, or more generally, towards the radical cation chain reactions occurring during pathogenesis. In this regard, the scavenging activity of CELLOX on short cellodextrins was higher than that of OGOX on short galacturonan oligomers (Table I), including a stronger laccase-counteracting activity (Fig. 4). Intriguingly, cellulose and lignin are strictly associated in lignocellulose^1^, suggesting that cellulose fragments with free-reducing ends could be exploited by CELLOX to preserve the integrity of lignocellulose by contrasting the radical cation-generating activity of microbial ligninases^16^. In the “OGOX1-OG” system, the scavenging efficiency of different FHS-OGOX1/OG-oligomer combinations towards ABTS^•+^ was inversely related to the length of each oxidizable oligomer, i.e., the shorter is the more scavenging (Table I), suggesting that these reaction mechanisms may differently contribute to plant defence depending on the progression of infection. For example, H_2_O_2_-dependent lignification can be mediated by extracellular plant PODs as soon as cell wall fragments are formed by microbial GHs and oxidized by plant OSOXs^9^, whereas the radical cation-scavenging activity is an additional effect occurring in a later phase, i.e., when cell wall oligosaccharides, including those already oxidized, are further converted into smaller oligomers by microbial GHs (Fig. 5b). Indeed, a strong scavenging activity in the first phase of infection could interfere with the radical-generating activity of plant peroxidases and other H_2_O_2_-mediated reactions (Fig. S3), making understandable why the activity of OGOX1 on degradation-intermediate products (e.g., longer oligomers such as OGs) was characterized by a negligible scavenging activity (Fig. S3, Fig. 4). In conclusion, our study, although limited to *in vitro* evidence, provides a novel perspective on how OSOXs can generate radical cation scavenging activity in the apoplast with a power proportional to the extent of degradation of plant cell wall, with possible implications for redox homeostasis and defense against oxidative stress during microbial infection. Indeed, the amount of H_2_O_2_ produced by OSOXs on short oligosaccharides was significantly reduced in the presence of a strong radical cation scavenging activity (Fig. S3-S4), suggesting a possible mechanism for modulating H_2_O_2_ homeostasis. Further studies will be required to understand whether the radical cation scavenging activity of OSOXs inhibits the production of H_2_O_2_ or consumes the H_2_O_2_ already present as obtained by the oxidation of short oligosaccharides. In this regard, it is noteworthy that millimolar concentrations of gluconolactone and H_2_O_2_ have been shown to non-enzymatically reduce Cr-(VI)^17^, indicating that a lactone-intermediate and H_2_O_2_ have the potential to synergistically act as reductants of (metal) cations. Knowledge of the enzymatic kinetics may be used to design further experiments to verify the occurrence of the reaction mechanisms described here also in an *in vivo* environment.

## METHODS

### Construction of the synthetic gene encoding FHS-OGOX1 by bioinformatic tools

The gene encoding the mature OGOX1.2 isoform from *Arabidopsis thaliana* (AT4G20830.2) was fused downstream of the SUMOstar sequence developed by LifeSensor Inc. (https://lifesensors.com/) that also included the sequences encoding the FLAG- (DYKDDDDK) and 6xHis-tags (HHHHHH) (https://lifesensors.com/wp-content/uploads/2019/09/2160_2161_Pichia_SUMOstar_Manual-1.pdf). The sequence of the chimeric gene, here referred to as *FHS-OGOX1*, was codon-optimized with the codon usage of *Pichia pastoris* by using the online tool OPTIMIZER (http://genomes.urv.es/OPTIMIZER/)^18^ and entirely synthesized by Genescript (https://www.genscript.com/) by adding the bases of the restriction sites PstI and XbaI at the 5^I^ and 3^I^ ends, respectively, of the gene. The gene was then cloned in pPICZαB expression vector (Invitrogen, San Diego, USA) in frame with the sequence encoding the yeast α factor for the secretion of recombinant proteins in the medium.

### Heterologous expression of FHS-OSOXs in *P. pastoris*

The recombinant FHS-CELLOX, i.e., a flag-his-sumoylated form of CELLOX from *A. thaliana* (AT4G20860), was expressed and purified as previously described^9^. The construct pPICZαB/FHS-OGOX1 was transformed in *E. coli* DH5α competent cells (ThermoFisher, Waltham, USA) for plasmid amplification. Then, the construct was linearized by SacI and introduced in *P. pastoris* by electroporation^19^. Multi-copy transformants were selected on solid YPDS medium [1% (w/v) yeast extract, 2% (w/v) peptone, 2% (w/v) dextrose, 1 M Sorbitol] with zeocin as antibiotic resistance marker (1 mg.mL^−1^). For protein expression, several colonies of multicopy *P. pastoris* transformants were inoculated in 5 mL of YPD medium supplemented with 200 μg.mL^−1^ zeocin and incubated at 28 °C in a rotary shaker at 180 rpm for 72-96h. The cultures were then centrifuged and the cell pellets resuspended in 1.5 mL of Buffered Minimal Medium [BMM; 0.1 M K-phosphate (pH 6.0), 1.34 % (w/v) YNB, 4 × 10-5 % (w/v) biotin and 0.5% (v/v) methanol] to induce the expression of FHS-OGOX1 and let them grow for additional 48 hours. To detect the expression of recombinant FHS-OGOX1, different culture filtrates were evaluated by SDS-PAGE/Coomassie blue staining. For FHS-OGOX1 purification, the methanol-induced culture filtrate from the highest expressing transformant was subjected to buffer exchange (50 mM Tris-HCl pH 7.5, 500 mM NaCl, 1 mM 2-mercaptoethanol and 10 mM imidazole) and then loaded onto a HisTrap HP column (ThermoFisher, Waltham, USA). The eluted fractions containing FHS-OGOX1 were pooled and dialyzed in 50 mM Tris-HCl pH 7.5, 100 mM NaCl and 100 mM (NH_4_)_2_SO_4_. Before its use in the assays, the purity of dialyzed FHS-OGOX1 preparation was evaluated by SDS-PAGE/Coomassie blue staining and quantified by UV-absorbance (ε280nm= 75440 M^−1^.cm^−1^). The activity of FHS-OGOX1 was evaluated by using the xylenol orange assay^6^. The activity of FHS-OGOX1 was assayed in 50 mM Na-Acetate pH 5.0 and 50 mM NaCl by using OGs, OG3 or OG4 (15 μM) as substrates in the presence of purified enzyme (4 nM), and then expressed as μmoles of H_2_O_2_ generated per minute per mg of FHS-OGOX1. The OG mixture (degree of polymerization: 10-15; average value used for mass-to-mole conversion: 2306 g.mol^−1^) and OG3 were purchased from Biosynth Carbosynth (https://www.carbosynth.com/) whereas OG4 was purchased from ELICYTIL (https://www.elicityl-oligotech.com/). For each galacturonan oligomer stock solution (1 mM), the amount of reducing ends was confirmed by the reducing sugar-assay^20^ using different amounts of galacturonic acid as calibration curve.

### Evaluation of FHS-OGOX1 activity by ABTS-HRP coupled and xylenol orange assays

Preliminarily, the OG-oxidizing activity of FHS-OGOX1 was measured by two different spectrophotometric methods, i.e., the ABTS-HRP coupled assay^8^ and the xylenol orange assay^6^. In both assays, the activity of FHS-OGOX1 was assayed in 50 mM Na-Acetate pH 5.0 and 50 mM NaCl by adding OGs or OG4 (15 μM) as substrates in the presence of the purified enzyme (4 nM). For the ABTS-HRP coupled assay, the absorbance at 415 nm from the enzymatic reactions was measured in continuous mode, subtracted to that of control reaction (basal mixture) and then converted into μmol ABTS^•+^ (ε415nm = 36 mM^−1^.cm^−1^). Conversion of μmol ABTS^•+^ into μmol H_2_O_2_ was obtained by applying the conversion factor [1.92 molecules ABTS^•+^: 1 molecule H_2_O_2_]^21^. Differently from the ABTS-HRP coupled assay, the xylenol orange assay is an end-point method^6^. Here, the absorbance at 560 nm from the same enzymatic reactions was measured at seven different time-points (0, 2, 4, 8, 16, 20 and 40 min), subtracted to that of control reaction (basal mixture) and converted into μmol H_2_O_2_ by interpolation with the H_2_O_2_-calibration curve. All the analysis were performed in triplicates at 25°C.

### Evaluation of scavenging activity of FHS-OSOXs by ABTS^•+^-reduction assay

In the ABTS^•+^-reduction assay, the starting concentration of ABTS (110 μM) was converted into 90 μM ABTS^•+^ and 20 μM ABTS by using K_2_S_2_O_8_ as ABTS-activating agent. Exhaustive K_2_S_2_O_8_-mediated oxidation of ABTS was achieved by incubating the reaction for 16 hours at 25°C. At the end of incubation, the amount of ABTS^•+^ was determined by absorbance at 415 nm (ε415nm = 36 mM^−1^.cm^−1^). The scavenging activity of FHS-OGOX1 towards ABTS^•+^ was evaluated in 50 mM Na-Acetate pH 5.0 and 50 mM NaCl, by adding the purified FHS-OGOX1 (4 nM) and OGs or the appropriate galacturonan oligomer (OG4, OG3) (15 μM). oxOG4 was prepared from OG4 as previously described^6^. The scavenging activity of FHS-CELLOX towards ABTS^•+^ was evaluated in 50 mM Na-Acetate pH 5.0 and 50 mM NaCl, by adding the purified FHS-CELLOX (16 nM) and the appropriate cellodextrin (CD4 or CD3) (15 μM). CD4 and CD3 were purchased from Megazyme (Bray, Ireland). For each cellodextrin stock solution (1 mM), the amount of reducing ends was confirmed by the reducing sugar-assay^20^ using different amounts of glucose as calibration curve. To evaluate the production of H_2_O_2_ in the ABTS^•+^-reduction assay, HRP VI-A type (P6782, Sigma-Aldrich, St. Louis, USA) was added to each reaction (1.25 μM). To determine the reduction of ABTS^•+^ over time, the absorbance at 415 nm from the enzymatic reactions was measured in continuous mode, subtracted to that of control reaction (basal mixture) and then converted into μmol ABTS^•+^ (ε415nm = 36 mM^−1^.cm^−1^). In parallel, the xylenol orange assay was used to measure the amount of H_2_O_2_ produced by FHS-OSOXs under the same reaction conditions [50 mM Na-Acetate pH 5.0 and 50 mM NaCl, pure FHS-OGOX1/CELLOX (4/16 nM) and the appropriate oligosaccharide (15 μM)]. All the analyses were performed in triplicates by using an Infinite^®^ M Nano200 spectrophotometer (Tecan AG, Männedorf, Switzerland). The scavenging efficiency was calculated as ratio between the net decrease of ABTS^•+^ and the production of H_2_O_2_ at a specific reaction time by considering that n moles of H_2_O_2_ corresponds to n moles of each oxidized oligomer. For the vary FHS-OSOX/oligomer combinations, the net decrease of ABTS^•+^ (decrease of ABTS^•+^ subtracted to that of the basal mixture with the only enzyme) was extrapolated at three different time-points from each reaction (4, 8 and 16 min), divided by the production of H_2_O_2_ at the same time-points and then averaged. For the FHS-OGOX1/OG3 combination, longer reaction times (8, 16 and 20 min) were considered since the absorbance values of shorter reaction times (4 min) were close to the lower limit of detection.

### Evaluation of ABTS-oxidizing activity of fungal laccase in the presence of FHS-OSOXs and short oligosaccharides

The ABTS-oxidizing activity of laccase from *Trametes versicolor* (38429, Sigma-Aldrich, St. Louis, USA) was evaluated in 50 mM Na-Acetate pH 5.0 and 50 mM NaCl, by using the ABTS-oxidation assay (100 μM ABTS) in the presence of the four FHS-OGOX1/OG4, FHS-OGOX1/OGs, FHS-CELLOX/CD3 and FHS-CELLOX/CD4 combinations. The activity was assayed by using the laccase alone (5 μg.mL^−1^) or in the presence of FHS-OGOX1 (4 nM) and OGs/OG4 (15 μM), or in the presence of FHS-CELLOX (16 nM) and CD3/CD4 (15 μM). For the ABTS-oxidation assay, the amount of ABTS^•+^ was spectrophotometrically measured at 25°C following the absorbance at 415 nm, subtracted to that of control reaction (basal mixture) and converted into μmoles of ABTS^•+^ (ε415nm = 36 mM^−1^.cm^−1^). In parallel, the xylenol orange assay was used to measure the amount of H_2_O_2_ produced by FHS-OSOXs in the presence of laccase (5 μg.mL^−1^) under the same reaction conditions [100 μM ABTS, 50 mM Na-Acetate pH 5.0 and 50 mM NaCl, pure FHS-OGOX1/CELLOX (4/16 nM) and the appropriate oligosaccharide (15 μM)]. All the analysis were performed in triplicates at 25°C.

## ACKNOWLEDGEMENTS

The authors gratefully acknowledge Prof. Felice Cervone, Prof. Giulia De Lorenzo and Dr. Daniela Pontiggia (Dept. of Biology and Biotechnology “C. Darwin”, University of Rome, Sapienza) for providing OGs, and Prof. Francesco Angelucci (Dept. of Life, Health and Environmental Sciences, University of L’Aquila) for inspiring discussion on flavoenzymes.

## AUTHOR CONTRIBUTIONS

M.B. and B.M. conceived the project. M.B. designed the experiments, A.S., V.S. and M.G. performed the experiments and analyzed the data jointly with M.B. and B.M.

A.S., V.S., M.G. and M.B. wrote the manuscript draft whereas M.B. and B.M. edited the final version of the manuscript. B.M. and M.B. supervised the research. All authors have approved the final manuscript.

## AVAILABILITY OF DATA

All relevant data are included in the article and/or its Supplementary Material. The datasets used and/or analyzed during the current study are available from M.B. on reasonable request.

## ADDITIONAL INFORMATION

Supplementary material (Fig. S1-S4) is provided as separated file.

## REFERENCES

1. Benedetti, M., Locci, F., Gramegna, G., Sestili, F. & Savatin, D. Green Production and Biotechnological Applications of Cell Wall Lytic Enzymes. Applied Sci. 9, 5012; 10.3390/app9235012 (2019).

2. Giovannoni, M., Gramegna, G., Benedetti, M. & Mattei, B. Industrial Use of Cell Wall Degrading Enzymes: The Fine Line Between Production Strategy and Economic Feasibility. Front Bioeng Biotechnol. 8, 356; 10.3389/fbioe.2020.00356 (2020).

3. Pontiggia, D., Benedetti, M., Costantini, S., De Lorenzo, G. & Cervone, F. Dampening the DAMPs: how plants maintain the homeostasis of cell wall molecular patterns and avoid hyper-immunity. Front Plant Sci. 11, 613259; 10.3389/fpls.2020.613259 (2020).

4. Voxeur, A., Habrylo, O., Guenin, S., et al. Oligogalacturonide production upon Arabidopsis thaliana-Botrytis cinerea interaction. Proc Natl Acad Sci USA. 116, 19743–19752 (2019).

5. Smirnoff, N. & Arnaud, D. Hydrogen peroxide metabolism and functions in plants. New Phytol. 221, 1197–1214 (2019).

6. Benedetti, M., Verrascina, I., Pontiggia, D., et al. Four Arabidopsis berberine bridge enzyme-like proteins are specific oxidases that inactivate the elicitor-active oligogalacturonides. Plant J. 94, 260–273 (2018).

7. Haddad Momeni, M., Fredslund, F., Bissaro, B., et al. Discovery of fungal oligosaccharide-oxidising flavo-enzymes with previously unknown substrates, redox-activity profiles and interplay with LPMOs. Nat Commun. 12, 2132; 10.1038/s41467-021-22372-0 (2021).

8. Scortica, A., Capone, M., Narzi, D., et al. A molecular dynamics-guided mutagenesis identifies two aspartic acid residues involved in the pH-dependent activity of OG-OXIDASE 1. Plant Physiol Biochem. 169, 171–182 (2021).

9. Scortica, A., Giovannoni, M., Scafati, V., et al. Berberine Bridge Enzyme-Like oligosaccharide oxidases act as enzymatic transducers between microbial glycoside hydrolases and plant peroxidases. Mol. Plant. Microbe Int. 10.1094/MPMI-05-22-0113-TA (2022).

10. Locci, F., Benedetti, M., Pontiggia, D., et al. An Arabidopsis berberine bridge enzyme-like protein specifically oxidizes cellulose oligomers and plays a role in immunity. Plant J. 98, 540–554 (2019).

11. Miller, N. J., Rice-Evans, C., Davies, M. J., Gopinathan, V. & Milner, A. A novel method for measuring antioxidant capacity and its application to monitoring the antioxidant status in premature neonates. Clin Sci (Lond). 84, 407–412 (1993).

12. Childs, R. & Bardsley, W. The steady-state kinetics of peroxidase with 2,2’-azino-di-(3-ethyl-benzthiazoline-6-sulphonic acid) as chromogen. Biochem. J. 145, 93–103 (1975).

13. Bourbonnais, R., Paice, M., Reid, I., Lanthier, P. & Yaguchi, M. Lignin oxidation by laccase isozymes from Trametes versicolor and role of the mediator 2,2’-azinobis(3-ethylbenzthiazoline-6-sulfonate) in kraft lignin depolymerization. Appl Environ Microbiol. 61, 1876–1880 (1995).

14. Hilgers, R., Vincke, J., Gruppen, H. & Kabel, M. Laccase/Mediator Systems: Their Reactivity toward Phenolic Lignin Structures. ACS Sustain Chem Eng. 6, 2037–2046 (2018).

15. Collins, P., Dobson, A. & Field, J. Reduction of the 2,2’-Azinobis(3-ethylbenzthiazoline-6-sulfonate) cation radical by physiological organic acids in the absence and presence of manganese. Appl Environ Microbiol. 64, 2026–2031 (1998).

16. Chen, H. Chemical composition and structure of natural lignocellulose in Biotechnology of Lignocellulose (ed. Springer, Dordrecht) 25–71 (2014).

17. Romo-Rodríguez, P., Acevedo-Aguilar, F. J., Lopez-Torres, A., Wrobel, K., Wrobel, K. & Gutiérrez-Corona, J. F. Cr(VI) reduction by gluconolactone and hydrogen peroxide, the reaction products of fungal glucose oxidase: Cooperative interaction with organic acids in the biotransformation of Cr(VI). Chemosphere. 134, 563–570 (2015).

18. Puigbò, P., Guzmán, E., Romeu, A. & Garcia-Vallvé, S. OPTIMIZER: a web server for optimizing the codon usage of DNA sequences. Nucleic Acids Res. 35, W126–131 (2007).

19. Wu, S. X. & Letchworth, G. J. High efficiency transformation by electroporation of Pichia pastoris pretreated with lithium acetate and dithiothreitol. BioTechniques. 36, 152–154 (2004).

20. Lever, M. A new reaction for colorimetric determination of carbohydrates. Anal Biochem. 47, 273–279 (1972).

21. Cai, H., Liu, X., Zou, J., et al. Multi-wavelength spectrophotometric determination of hydrogen peroxide in water with peroxidase-catalyzed oxidation of ABTS. Chemosphere. 193, 833–839 (2018).

