## Supplementary Material for "RADICAL CATION SCAVENGING ACTIVITY OF BERBERINE BRIDGE ENZYME-LIKE OLIGOSACCHARIDE OXIDASES ACTING ON SHORT CELL WALL FRAGMENTS"

### TABLE OF CONTENTS

#### Supplementary Figures (Fig. S1-S4)

Figure S1. Heterologous expression of FHS-OGOX1 in *P. pastoris*.

Figure S2. Reduction of $ABTS^{\bullet+}$ requires the activity of FHS-OGOX1 on OG4.

Figure S3. Reduction of $ABTS^{\bullet+}$ by the activity of FHS-OGOX1 on different OG-oligomers in the presence of HRP.

Figure S4. Production of $H_2O_2$ by the activity of different OSOX/oligomer combinations in the presence of laccase.

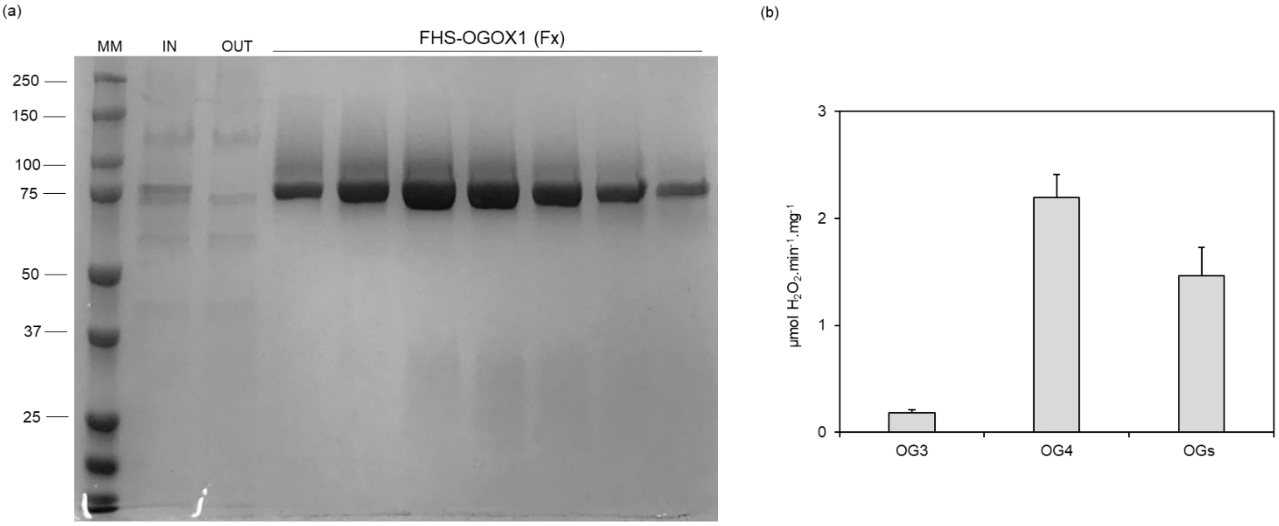


**Figure S1. Heterologous expression of FHS-OGOX1 in *P. pastoris*.** (a) SDS-PAGE/Coomassie blue staining analysis of different fractions (Fx) containing FHS-OGOX1 as eluted from the his-trap column. The sample before (IN) and after (OUT) the passage through the column was also analyzed. Molecular weight marker (MM) is also reported. (b) Oxidizing activity of FHS-OGOX1 (µmol $H_2O_2$.min$^{-1}$.mg$^{-1}$) at pH 5.0 using OG-oligomers with different length as substrates. Values are mean ± SD (N=3). [FHS-OGOX1: flag-his-sumoylated oligogalacturonide-oxidase 1, OGs: oligogalacturonides, OG4: tetra-galacturonic acid, OG3: tri-galacturonic acid].

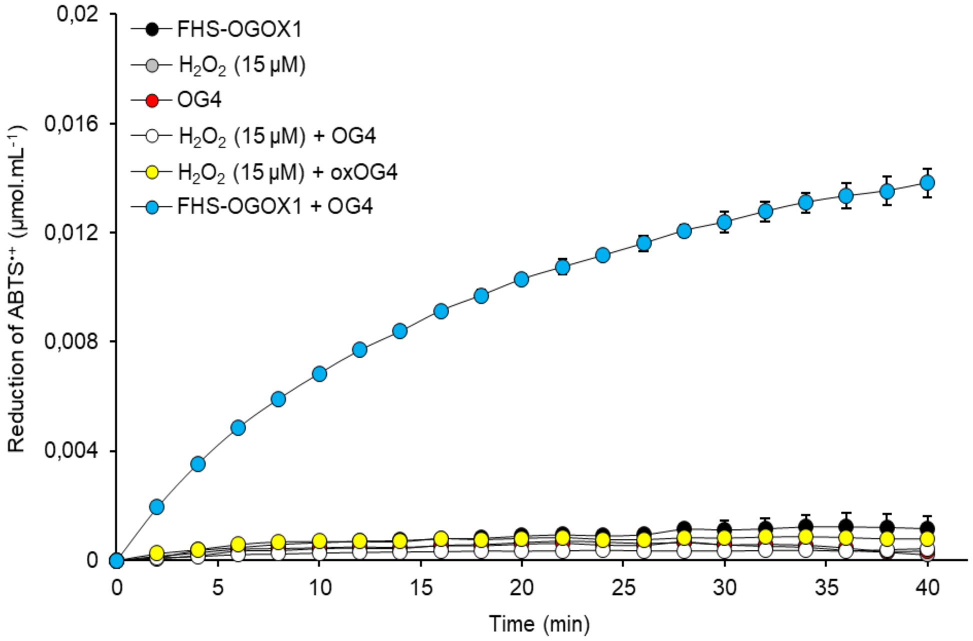


**Figure S2. Reduction of ABTS$^{\bullet+}$ requires the activity of FHS-OGOX1 on OG4.** Reduction of ABTS$^{\bullet+}$ (µmol.mL$^{-1}$) over time at pH 5.0 using different combinations of substrates and reactants. Values are mean ± SD (N=3). [FHS-OGOX1: flag-his-sumoylated oligogalacturonide-oxidase 1, OG4: tetra-galacturonic acid, oxOG4: oxidized tetra-galacturonic acid].

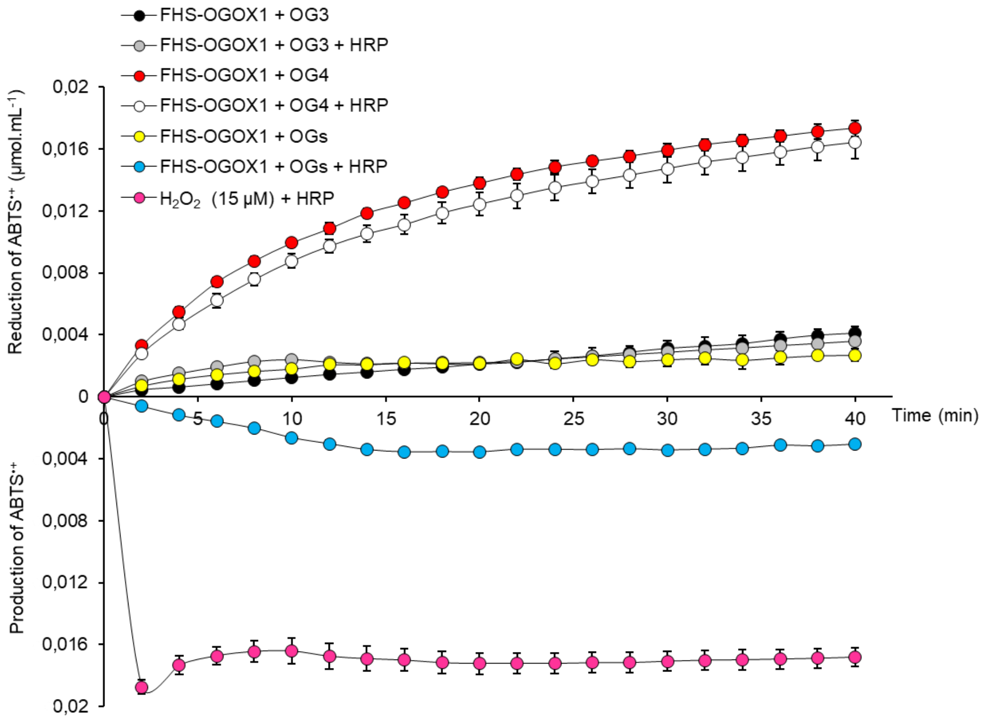


**Figure S3. Reduction of ABTS$^{•+}$ by the activity of FHS-OGOX1 on different OG-oligomers in the presence of HRP.** Reduction of ABTS$^{•+}$ (µmol.mL$^{-1}$) over time at pH 5.0 by the activity of FHS-OGOX1/OG3, FHS-OGOX1/OG4 and FHS-OGOX1/OGs combinations in the presence of HRP (+ HRP). [$H_2O_2$ + HRP] is reported as positive control of HRP-mediated oxidation of ABTS. Values are mean ± SD (N=3). [FHS-OGOX1: flag-his-sumoylated oligogalacturonide-oxidase 1, HRP: horseradish peroxidase VI-type, OGs: oligogalacturonides, OG4: tetra-galacturonic acid, OG3: tri-galacturonic acid].

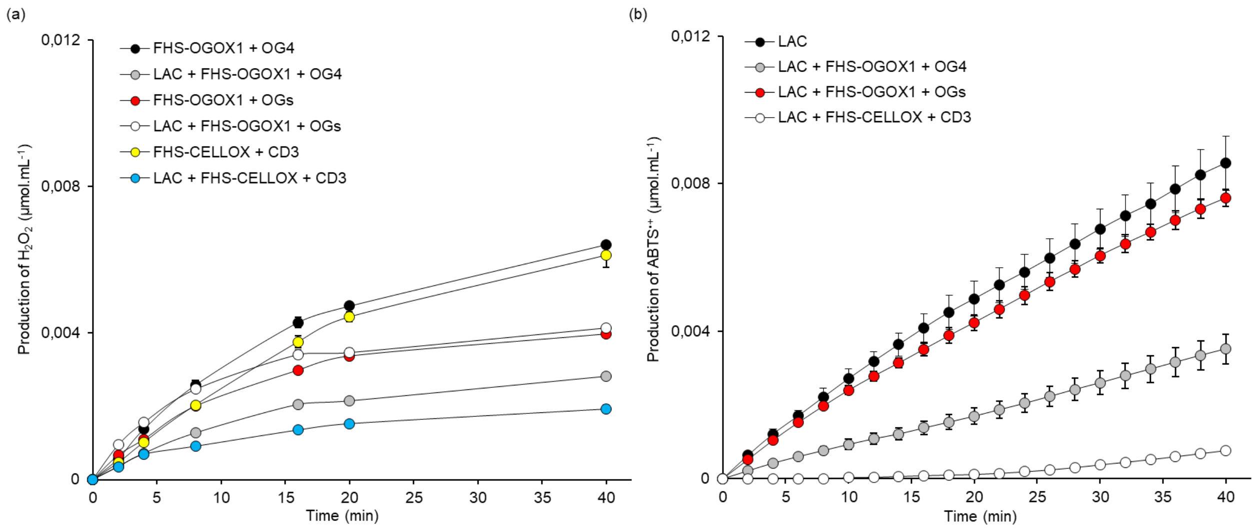


**Figure S4. Production of $H_2O_2$ by the activity of different OSOX/oligomer combinations in the presence of laccase.** (a) Production of $H_2O_2$ (µmol.mL$^{-1}$) over time at pH 5.0 by the activity of FHS-OGOX1/OG4, FHS-OGOX1/OGs and FHS-CELLOX/CD3 combinations in the presence of laccase as determined by the xylenol orange assay. (b) Production of ABTS$^{\bullet+}$ over time by laccase alone and in the presence of the same OSOX/oligomer combinations shown in (a) (extrapolated from Fig. 4). Values are mean ± SD (N=3). [CD3: cellotriose, FHS-CELLOX: flag-his-sumoylated cellodextrin-oxidase, FHS-OGOX1: flag-his-sumoylated oligogalacturonide-oxidase 1, LAC: laccase from *T. versicolor*, OGs: oligogalacturonides, OG4: tetra-galacturonic acid].
